# A novel approach to identify cross-identity peptides between Epstein-Barr virus and central nervous system proteins in Guillain-Barré syndrome and multiple sclerosis

**DOI:** 10.1101/2023.10.17.562729

**Authors:** Helmut Kennedy Azevedo do Patrocínio, Tayná da Silva Fiúza, Jonas Ivan Nobre Oliveira, João Firmino Rodrigues-Neto, Selma Maria Bezerra Jerônimo, Gustavo Antônio de Souza, João Paulo Matos Santos Lima

## Abstract

**Background:** Guillain-Barré Syndrome (GBS) and multiple sclerosis are autoimmune diseases associated with an immune system attack response against peripheral and central nervous system autoantigens, respectively. Given the potential of Epstein-Barr virus (EBV) as a risk factor for both multiple sclerosis and GBS, the present study aimed to identify crucial residues among potential EBV CD4+ T lymphocyte epitopes and nervous system proteins.

**Methods:** Public databases (Allele Frequency Net Database, Immune Epitope Database, Genevestigator and Protein Atlas) were used to select proteins abundant in the nervous system, EBV immunogenic proteins, and HLA haplotypes. Computational tools were employed for predicting HLA-binding peptides and immunogenicity. For this, we developed immuno-cross, a Python tool (https://github.com/evoMOL-Lab/immuno-cross) to compare residue identity among nonamers.

**Results:** We found ten proteins from the nervous system and 28 from EBV, which were used for predicting the binding peptides of 21 common HLAs in the world population. A total of 1411 haplotypes were distributed among 51 pairs of HLAs. Simulations were performed to determine whether nonamers from the EBV and nervous system proteins targeted TCR-contact residues. Then, three selection criteria were used, based on the relevance of each contact in the TCR-peptide-MHC interaction. The primary contact has to be located at position P5, and the positions P2, P3, and P8 were weighed as secondary, and P4, P6, and P7 were considered tertiary. Nonamers of EBV proteins and myelin proteins were combined in pairs and compared based on predefined selection criteria. The Periaxin protein had the highest number of nonamers pairs among PNS proteins, with 35 pairs. Four nonamers pairs from APLP1, two from CNP, and two from MBP bind to alleles of the haplotype DR-15.

**Conclusions:** The new approach proposed herein revealed that peptides derived from nervous system and EBV proteins share identical residues at critical contact points, which supports molecular mimicry. These findings suggest cross-reactivity between them and that the nonamer pairs identified with this approach have the potential to be an autoantigen. Experimental studies are needed to validate these findings.

## Introduction

Guillain-Barré syndrome (GBS) is an acute neuropathy characterized by an autoimmune response against the peripheral nervous system (PNS) (1). GBS is the most common cause of acute flaccid paralysis, after the control of poliomyelitis, with an average global prevalence of up to 1.9 cases per 100.000 person-year (2). GBS recovery phase may last from weeks to years (3). History of Post Infections or vaccinations are commonly associated with onset of GBS. This disease has a broad spectrum that includes two main subtypes: Acute Motor Axonal Neuropathy and Acute Inflammatory Demyelinating Polyradiculoneuropathy (AIDP). *Campylobacter jejuni* infection is highly associated with triggers one (3), whereas there is no established pathological mechanism for triggering AIDP. Some studies show a higher frequency of viral infections before the onset of AIDP (4–6), though not yet associated with a known autoimmune antigen. As an autoimmune disease, losing immune tolerance to self-antigens may play a vital role in developing GBS (3,7).

Multiple sclerosis (MS) is a nervous system-afflicting autoimmune disease, atargenting central nervous system (CNS) cells, causing focal areas of chronic inflammation and demyelination of white and gray matter, and spinal cord (8). MS global prevalence is about 36 per 100,000 population (9). Similar to GBS, epidemiological studies suggest that an Epstein-Barr virus (EBV) infection may be associated with MS, although its role in the pathology remains unclear (10). Anti-EBNA1 (Epstein–Barr virus nuclear antigen 1) antibody titers can be detected in approximately 99.5% of patients with MS before the onset of clinical symptoms, in contrast to 94% in healthy subjects (11). Furthermore, a prior EBV infection has significantly increased the risk of developing MS by 32-fold, potentially rendering it the primary risk factor for the disease (11). A possible mechanism that could explain how EBV triggers MS is molecular mimicry between viral and CNS antigens (12,13).

As EBV may pose a potential risk factor for both MS and the demyelinating form of GBS, the present investigation aims to identify crucial residues among potential epitopes of CD4+ T lymphocytes of EBV and nervous system proteins as a means to suggest how infection by this virus can contribute to the pathogenesis of diseases. The work presented here employs immunoinformatics tools to generate binding core affinity predictions. It uses a novel binding-core identity analysis to rank crucial residues and peptides in solid binders. To the best of our knowledge, previous works have also examined the potential for molecular mimicry based on nonamer identity but have yet to consider the relevance of each position for TCR recognition. The three identity-based nonamer selection rules defined in this work aid in selecting TCR-context-dependent peptides, and our findings may guide wet lab experiments to reduce epitope test datasets.

## Material and Methods

We retrieved the primary data for this study from two distinct datasets: Protein Atlas (https://www.proteinatlas.org/) and Immune Epitope Database (IEDB) (www.iedb.org). We used the list Greenbaum et al. (14) proposed for global common HLA alleles and retrieved their frequency from the Allele Frequency Net Database (AFND). Figure 1 summarizes the experimental approach used.

**Figure 1:**
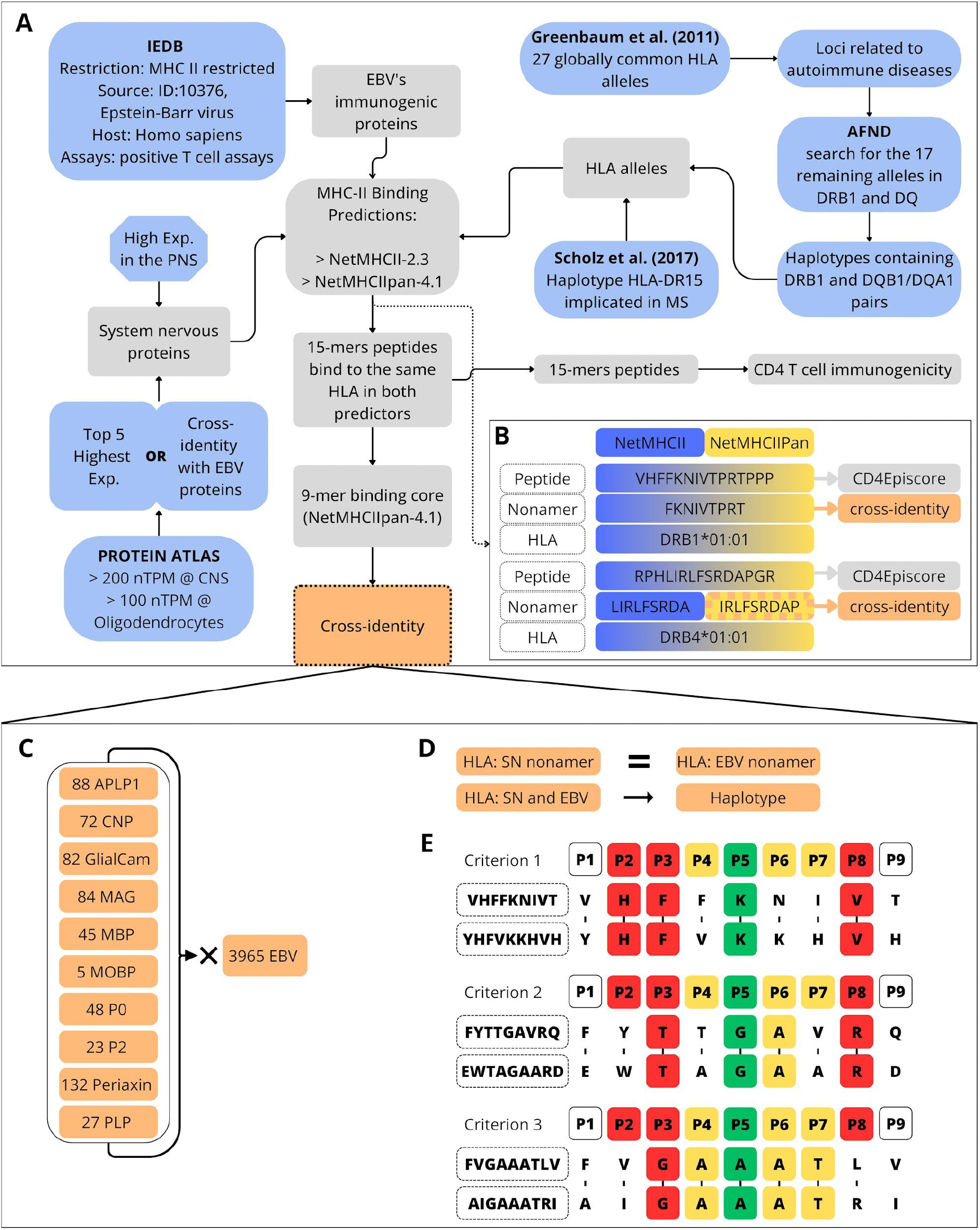
Scheme describing the methodology workflow to identify cross-identity peptides. **A**. Data retrieving steps (blue), input data, and subsequent analysis (gray). **B**. We selected 15-mer peptides predicted by both methods and for the same HLA. After, as nonamers may differ between methods, we chose the predictions from NetMHCIIpan. The 15-mer peptides were analyzed using the CD4 episcore, and the nonamers were assessed for cross-identity. **C**. The search for cross-identity of the peptides from human nervous system proteins against peptides from Epstein-Barr virus. **D**. The nonamers should bind to the same HLA or HLAs present in haplotype. **E**. The pipeline uses three rules for selecting nonamers with relevant identities interacting with the TCR. It starts with a minimal identity of 44% between the residues of two given nonamers and identical residues at position 5 (P5) as base criteria and the cross-identity of nonamers that bind to the same HLA allele or HLA alleles contained in a common haplotype. The three additional rules used to select nonamers where TCR-relevant cross-identity may occur. Criterion 1: identical residues at P2, P3, and P8 (P2-P3-P5-P8). Criterion 2: two identical residues at P2, P3 or P8, and at least one identical residue at P4, P6 or P7 (P5 - 2 x P2/P3/P8 - 1 x P4/P6/P7). Criterion 3: identical residues at P4, P6 and P7 and at least one identical residue at P2, P3 or P8 (P5 - 1 x P2/P3/P8 -P4-P6-P7).

### Selection of proteins from the human nervous system

Proteins were retrieved from the Protein Atlas data bank (15), using filters to select highly expressed proteins in the Central Nervous System (>200 nTPM) and in oligodendrocytes (>100 nTPM) on [June 1, 2022]. The supplementary material lists the entire query employed. Then, we performed a detailed investigation of the remaining entities using the Protein Atlas page for each protein to remove those that were also highly expressed in other tissues. We also used Genevestigator (https://genevisible.com/search) (16) to ensure that the selected proteins have high expression rates in the Central Nervous System. Since the Protein Atlas database has no specific parameters for the Peripheral Nervous System, we retrieved proteins from the Peripheral Nervous System proteomes and then reverse engineered those that may be the origin of GBS autoantigens, following confirmation of their expression using Genevestigator. The following analysis steps involved the selection of the top five highly expressed proteins from the brain tissue. We also included five other proteins in this interest group: two for the evidence suggesting their cross-identity or immunogenicity in either EBV or MS and three for their abundance in the PNS.

#### Frequent Haplotype Selection

To select HLA haplotypes relevant to this study, 27 globally common HLA alleles (14) were filtered out, considering only 17 of the DRB1 and DQ loci, mainly related to autoimmune diseases (17). The remaining alleles were then submitted to the AFND (http://www.allelefrequencies.net) (18) for the haplotype search. Haplotypes containing DRB1 and DQB1/DQA1 pairs were selected in the Haplotype Frequency Search of AFND (http://www.allelefrequencies.net/hla6003a.asp) using one of the scripts from Immuno-cross (https://github.com/evoMOL-Lab/immuno-cross) for collecting and processing the data. The final haplotype collection also contained a single external haplotype - HLA-DR15 [DRB1*15:01 (DR2b) and DRB5*01:01 (DR2a)], added for its implications in MS (19).

#### Epstein-Barr Immunogenic Proteins

Epstein-Barr proteins capable of eliciting CD4+ T lymphocyte responses were retrieved from the curated IEDB (20) querying epitopes (a) restricted to MHC class II, (b) with *Human herpesvirus 4 (Epstein Barr virus) (ID:10376, Epstein-Barr virus)* as the source organism, (c) *Homo sapiens* as host species and (d) positive for T lymphocyte assays. We then retrieved the entire sequence of the resulting EBV (strains Raji, GD1, B95-8, Cao, AG876) proteins from Uniprot (21).

### Peptide-HLA binding prediction

Using two methods, NetMHCII-2.3 (22) and NetMHCIIPan-4.1 (23), we assessed the capacity of peptides from immunogenic EBV proteins and nervous system proteins to bind to specific HLA alleles. The peptide’s length was 15 amino acids long. All peptides classified as ligands, whether strong or weak, were selected. We cross-referenced the results from the two predictors to identify the 15-mers peptides likely to bind to both methods and to the same HLA (Figure 1C). The predictions also indicate the core nonamers, which are peptides with nine residues, which constitute the primary component of the T cell epitope (24). Nonamers predicted by both methods may be the same, but, in the case they differ, we selected those predicted by NetMHCIIpan-4.1, known for its superior performance (23).

### EBV and Nervous System nonamer cross-identity

We compared the immunogenic nonamers from both groups considering residues in critical positions for T-cell receptor (TCR) interaction using another script included in our tool. Cross-reactivity does not require that all residues of the nonamer to be the same but that there is identity in some critical contacts with the TCR (25–30). Therefore, we formulated three criteria for selecting nonamers with relevant identities interacting with the TCR (Figure 1D, E).

Starting from a minimal identity of 44% (4 out of 9) between the residues of two given nonamers, and identical residues at position 5 (P5) as base criteria, additional rules were defined to select nonamers where TCR-relevant cross identity may occur: a) identical residues at P2, P3 and P8 (P2-P3-P5-P8); or b) two identical residues at P2, P3 or P8, and at least one identical residue at P4, P6 or P7 (P5 - 2 x P2/P3/P8 - 1 x P4/P6/P7); or c) identical residues at P4, P6 and P7 and at least one identical residue at P2, P3 or P8 (P5 - 1 x P2/P3/P8-P4-P6-P7). These criteria are only valid for evaluating the cross-identity of nonamers that bind to the same HLA allele or HLA alleles contained in a common haplotype. As an additional metric for evaluating the proximity of two nonamers, we used a similarity score that considers the substitution scores from residues 2 to 8 according to the BLOSUM62 matrix. We also used a script that implements the modules *substitution_matrices* and *pairwise2* of the Biopython package (version: 1.79) (31).

### CD4 T-cell immunogenicity prediction

The 15-mer peptides were submitted to the CD4episcore server (http://tools.iedb.org/CD4episcore/) to assess their ability to stimulate the CD4 T-lymphocyte response (32,33). The threshold classification of the peptides was 50%. The score combines the immunogenicity score and the 7-allele method score.

### Data processing and Visualization

We joined all the post-processing scripts, written in Python (v 3.10.4) to clean, filter, and extract data in the tool *Immuno-cross*, available at https://github.com/evoMOL-Lab/immuno-cross. We used the packages matplotlib (https://matplotlib.org/) (version 3.5) and seaborn (https://seaborn.pydata.org/) (version 0.11.2) to build the graphs. For visualizing the identity analysis of the nonamers, we used the package UpSetPlot (https://upsetplot.readthedocs.io/en/stable/) (v. 0.6.1).

## Results

### Protein selection and retrieving

#### Nervous System Proteins

The five proteins selected from the Protein Atlas (nPTM threshold) and validated by Genevisible as brain-specific high-expression were: Myelin basic protein (MBP), PLP1, 2’,3’-cyclic nucleotide 3’ phosphodiesterase (CNP), Amyloid beta precursor-like protein 1 (APLP1) and Myelin associated oligodendrocyte basic protein (MOBP). We also selected Two additional proteins, Hepatic and glial cell adhesion molecule (HepaCam) and myelin-associated glycoprotein (MAG), due to their implication in cross-reactivity in multiple sclerosis (12,34). Table 1 describes their nPTM and reference identifiers. Three abundant proteins in the PNS were selected: Myelin protein P0, Myelin protein P2, and Periaxin (35,36).

**Table 1.** Summary of nervous system proteins selected for peptide-HLA binding prediction.

| Protein | Abbreviation | Brain Expression (nPTM) | Oligodendrocytes Expression (nPTM) | Protein atlas | Genevisible |
| --- | --- | --- | --- | --- | --- |
| Myelin basic protein | MBP | 32454.4 | 2895.5 | ENSG00000197971 | P02686 |
| Proteolipid protein 1 | PLP1 | 7049.5 | 4822.5 | ENSG00000123560 | P60201 |
| 2',3'-cyclic nucleotide 3'phosphodiesterase | CNP | 1435.7 | 555.6 | ENSG00000173786 | P09543 |
| Amyloid beta precursor like protein 1 | APLP1 | 1182.3 | 461.3 | ENSG00000105290 | P51693 |
| Myelin associated oligodendrocyte basic protein | MOBP | 933.7 | 1607 | ENSG00000168314 | Q13875 |
| Excitatory amino acid transporter 1 | SLC1A3 | 783.1 | 249 | ENSG00000079215 | P43003 |
| Myelin associated glycoprotein | MAG | 690 | 267.8 | ENSG00000105695 | P20916 |
| Ermin | ERMN | 533.8 | 290.9 | ENSG00000136541 | Q8TAM6 |
| Neurotrimin | NTM | 324.3 | 1257.6 | ENSG00000182667 | Q9P121 |
| Hyaluronan and proteoglycanlink protein 2 | HAPLN2 | 276.4 | 202.1 | ENSG00000132702 | Q9GZV7 |
| Serpin family I member 1 | SERPINI1 | 266.2 | 174.7 | ENSG00000163536 | Q99574 |
| Formin like 2 | FMNL2 | 233 | 2442 | ENSG00000157827 | Q96PY5 |
| Hepatic and glial cell adhesion molecule | HEPACAM | 229.3 | 163.2 | ENSG00000165478 | Q14CZ8 |
| Myelin protein P0 | MPZ |  |  | ENSG00000158887 | P25189 |
| Periaxin | PRX |  |  | ENSG00000105227 | Q9BXM0 |
| Myelin protein P2 | PMP2 |  |  | ENSG00000147588 | P02689 |

Yellow, proteins selected by brain expression nPTM threshold; Green, additional proteins selected; Purple, Abundant proteins selected from the PNS.

#### Epstein-Barr Immunogenic Proteins

The 28 immunogenic proteins retrieved at IEDB contain 208 epitopes validated in 632 assays and contain envelope glycoproteins, nuclear antigens as most common in the group. Table 2 shows the number of epitopes and validating assays for each.

**Table 2.** Summary of Epstein-Barr proteins selected for peptide-HLA binding prediction.

| Protein | Abbreviation | Epitopes | Assays |
| --- | --- | --- | --- |
| Apoptosis regulator BHRF1 | BHRF1 | 7 | 12 |
| DNA polymerase catalytic subunit | BALF5 | 1 | 22 |
| DNA polymerase processivity factor BMRF1 | BMRF1 | 1 | 4 |
| Envelope glycoprotein B | BALF4 | 9 | 14 |
| Envelope glycoprotein GP350 | BLLF1 | 3 | 8 |
| Envelope glycoprotein H | BXLF2 | 1 | 6 |
| Envelope glycoprotein L | BKRF2 | 1 | 1 |
| Epstein-Barr nuclear antigen 1 | EBNA1 | 68 | 230 |
| Epstein-Barr nuclear antigen 2 | EBNA2 | 8 | 51 |
| Epstein-Barr nuclear antigen 3 | EBNA3 | 3 | 10 |
| Epstein-Barr nuclear antigen 4 | EBNA4 | 4 | 4 |
| Epstein-Barr nuclear antigen 6 | EBNA6 | 16 | 50 |
| Epstein-Barr nuclear antigen leader protein | EBNA-LP | 1 | 1 |
| Glycoprotein 42 | BZLF2 | 1 | 1 |
| Inner tegument protein | BOLF1 | 1 | 1 |
| Large tegument protein deneddylase | BPLF1 | 2 | 5 |
| Latent membrane protein 1 | LMP1 | 15 | 53 |
| Latent membrane protein 2 | LMP2 | 14 | 70 |
| Major capsid protein | MCP | 5 | 15 |
| Major DNA-binding protein | BALF2 | 2 | 4 |
| Major tegument protein | BNRF1 | 18 | 20 |
| mRNA export factor ICP27 homolog | BMLF1 | 1 | 5 |
| Nuclear egress protein 2 | NEC2 | 1 | 2 |
| Replication and transcription activator | BRLF1 | 1 | 2 |
| Ribonucleoside-diphosphate reductase small subunit | RIR2 | 1 | 3 |
| Secreted protein BARF1 | BARF1 | 17 | 17 |
| Trans-activator protein BZLF1 | BZLF1 | 4 | 11 |
| Triplex capsid protein 1 | TRX1 | 2 | 10 |

#### Haplotype Search and Allele Selection

With the resulting 21 HLA alleles (four HLAs DRB3/4/5, six DQA1/DQB1, and eleven DRB1), 66 haplotype pairs were obtained by combining DRB1 and DQA1/DQB1. Fifty-one were available in AFND, while 15 were not yet registered. 19 of the 51 haplotypes have reported frequencies equal to or higher than 1%, and the remaining 32 rare haplotypes (frequency < 1%) were discarded (Table S1, figure 2). In total, we located 1411 haplotypes distributed among 51 pairs of HLAs.

**Figure 2.**
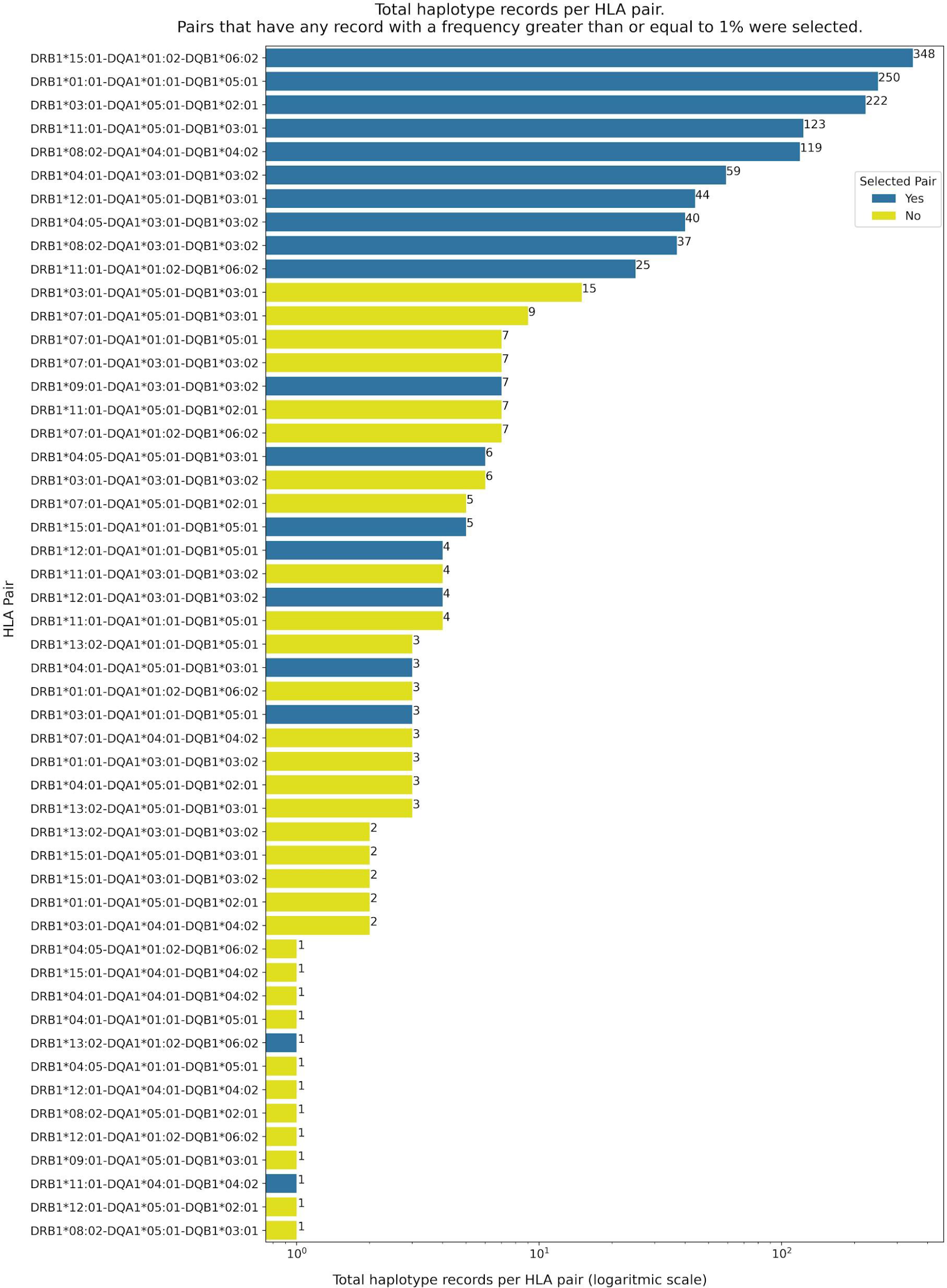
- HLA haplotypes obtained from AFND selected for the prediction of peptides. Blue: selected haplotypes pairs. Yellow: haplotypes not selected.

### Peptide-HLA binding prediction

The prediction of HLA ligands for nervous system proteins resulted in 5904 peptides by the NetMHCII and 4206 by NetMHCIIPan tools. The prediction of EBV peptides generated 86325 peptides with NetMHCII and 52944 with NetMHCIIPan. After filtering and identifying peptides predicted for both servers and ligands to the same HLA, the final result has 1695 peptides for nervous system proteins and 24912 for EBV proteins. Figure 3 shows the result for each of the nervous system proteins and the EBV proteins.

**Figure 3.**
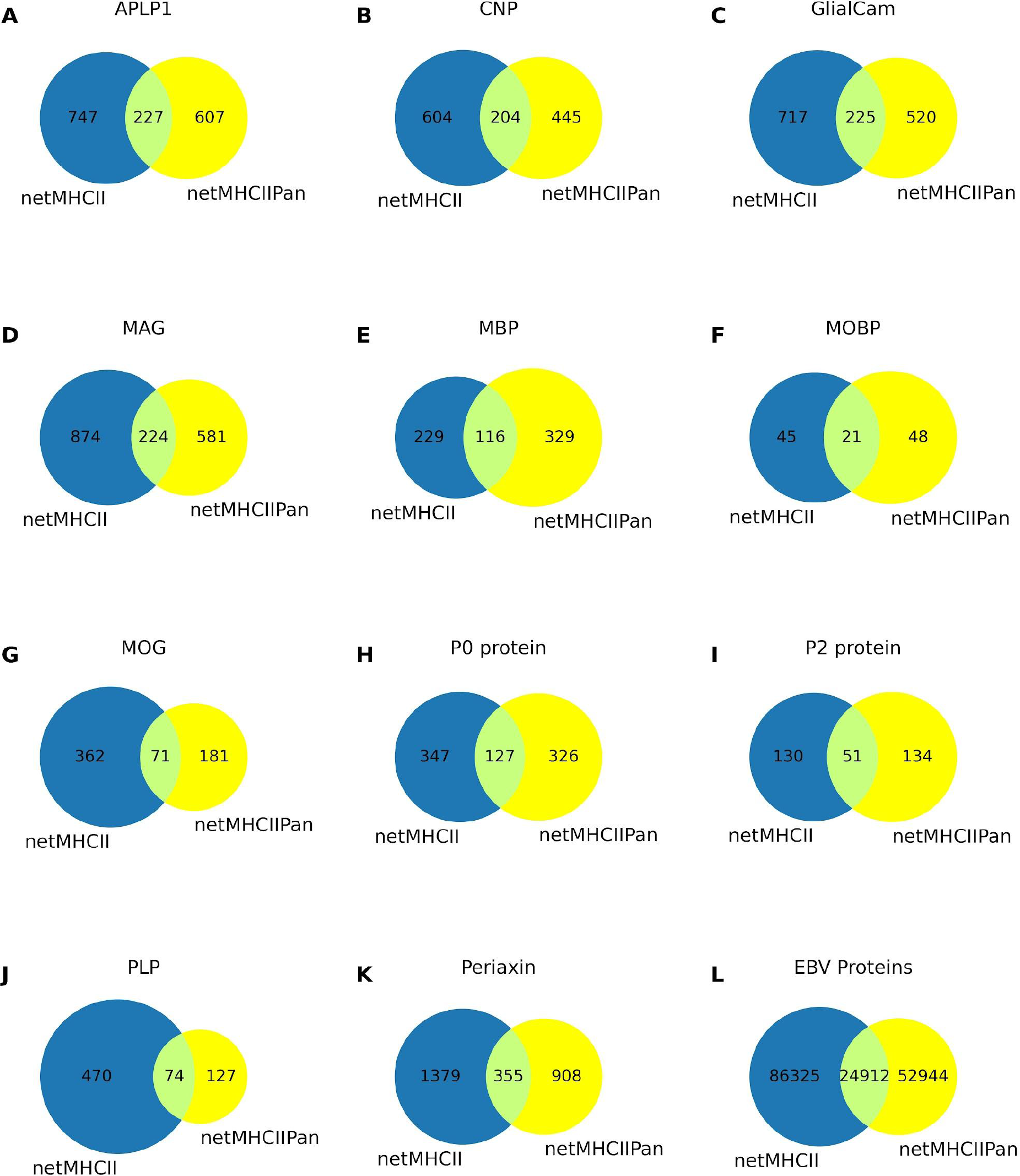
- Amount of ligands predicted by NetMHCII and NetMHCIIPan servers and by both for each nervous system protein and EBV proteins. Blue: total binders predicted by NetMHCII. Yellow: total binders predicted by NetMHCIIPan.

### EBV and Nervous System nonamer cross-identity and immunogenicity prediction

We compared nonamers of EBV strains to nervous system nanomers predicted to bind to the selected HLAs. The results of the nonamer cross-identity of the APLP1, CNP, GlialCam, MAG, MBP, MOBP, P0, P2, Periaxin, and PLP proteins are shown in tables S2–S11, respectively. Figure 4 summarizes the results for all proteins. We highlighted differences in the number of pairs selected by each selection criterion. All proteins have a predominance of pairs selected according to criterion 2 (P5 - 2 x P2/P3/P8 - 1 x P4/P6/P7), probably because this contains the highest number of combinations of identical residues. Since Criterion 3 (P5 - 1 x P2/P3/P8 - P4-P6-P7) only allows pairs of five or more identical residues and has few combinations, it has the lowest number of nonamers. Only three proteins complied with this criterion: APLP1, Periaxin, and **PLP**. Only APLP1, MBP, Periaxin, GlialCam, and PLP proteins showed peptides with immunogenic potential (Table 3).

**Figura 4.**
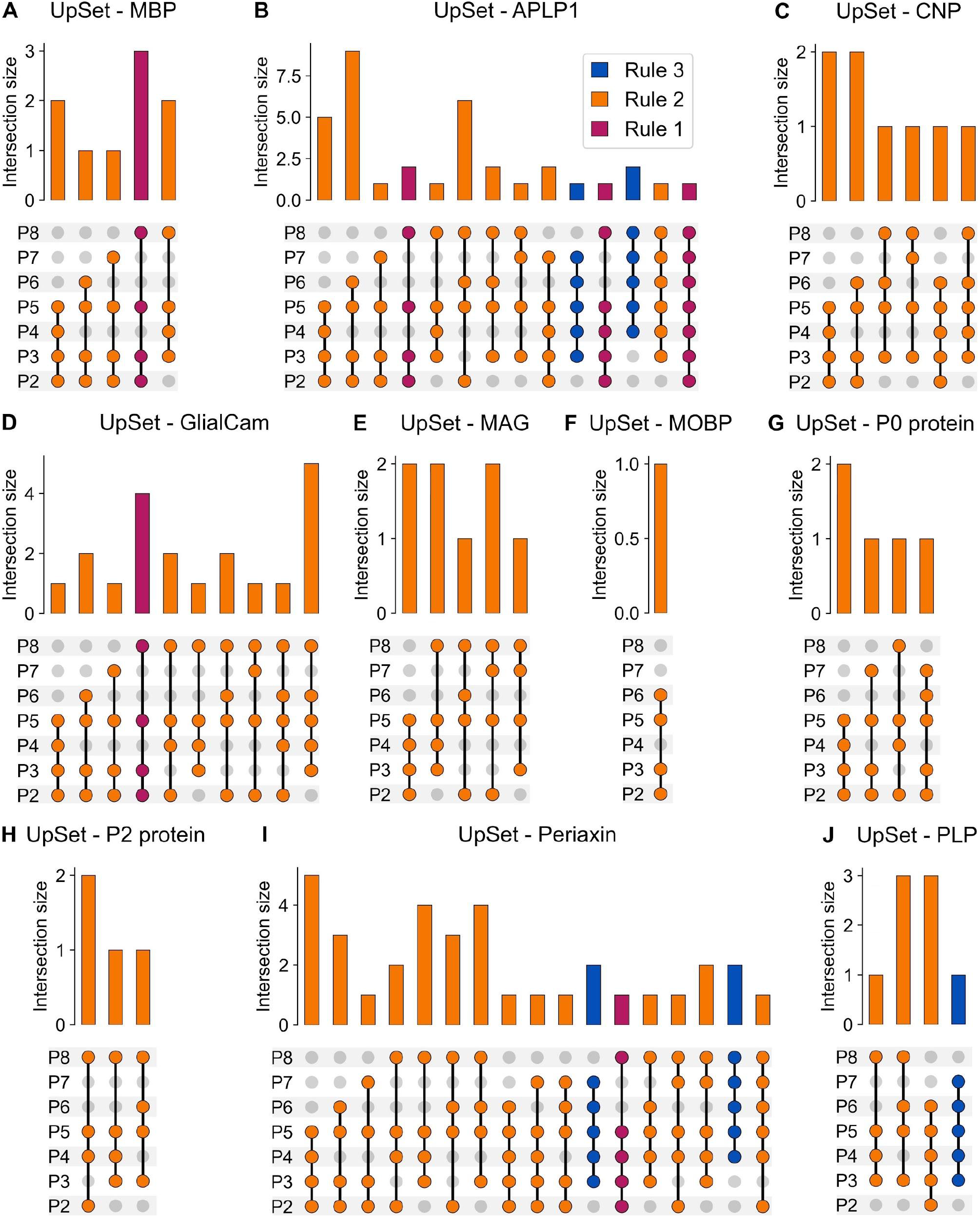
- UpSet plots summary of cross-identity between nonapeptides of nervous system proteins and EBV. We categorized the selected peptides according to the rule they complied with and grouped them according to the positions that presented identity. The "Intersection size" graph shows the size of each set. Purple: sets of peptides selected by rule 1. Orange: sets of peptides selected by rule 2. Blue: sets of peptides selected by rule 3.

**Table 3.**
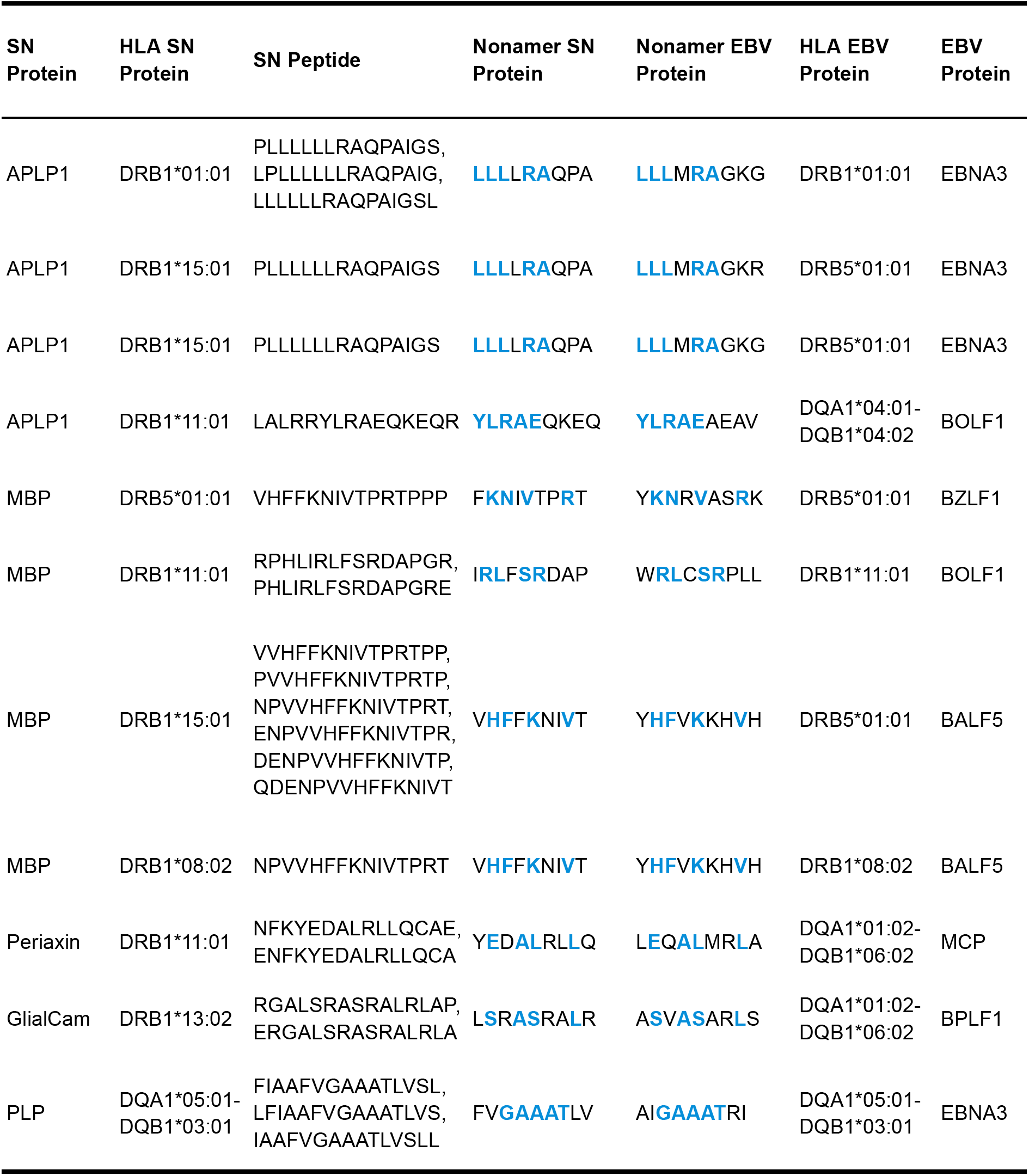
15-mer peptides immunogenicity prediction.

#### Amyloid Beta Precursor Like Protein 1 (APLP1)

We found that 35 nonamers from the APLP1 protein had a relevant similarity with EBV nonamers (Table S2, figure 4B). The HLA-DQA1*05:01/DQB1*03:01 stands out because it binds to 11 pairs of nonamers. The pair formed by the VEGAEDEEE^APLP1 and AEGAEDEEG^BOLF1 displayed the highest similarity score (36) and identity (7/9) among all proteins. This APLP1 nonamer is also present in another five pairs, three of which were selected by criterion 1 (P2-P3-P5-P8). The other pair selected by criterion 1 (P2-P3-P5-P8) was MLTLEEQQL^APLP1/SLTLEPIQD^BMLF1, with five identical residues. Four pairs (LLLLRAQPA^APLP1/LLLMRAGKG^EBV, LLLLRAQPA^APLP1/LLLMRAGKG^EBV, YLRAEQKEQ^APLP1/YLRAEAEAK^EBV and NQSLGLLDQ^APLP1/MQSYGLERL^EBV) bind to alleles of the DR15 haplotype (DRB1*15:01, DRB5*01:01, and DQA1*01:02-DQB1*06:02), which is a known risk factor in MS (37). The nonamers LLLLRAQPA and YLRAEQKEQ stand out because they occur in 15-mer peptides predicted to be immunogenic (Table 3).

#### Hepatic And Glial Cell Adhesion Molecule (GlialCam)

Twenty pairs of GlialCam protein nonamers were selected (Table S4, figure 4D). Five pairs, all with five identical residues, are formed by the nonamers LVASTTVLE from GlialCam and YEASTTYLS from the EBV BXLF2 protein because both bind to HLAs DRB1*04:05, DQA1*03:01-DQB1*03:02, DRB1*04:01, DRB1*09:01 and DRB1*13:02, presented in haplotypes. The YSVSPAVPG nonamer forms five pairs, four selected by criterion 1 (P2-P3-P5-P8). One nonamer, LSRASRALR, is present in 15-mer peptides predicted to be immunogenic (Table 3). The pair ERGALSRAS^GlialCam/SEGALALAG^EBV binds to HLA-DQA1*01:02-DQB1*06:02.

#### Myelin Basic Protein (MBP)

The MBP protein had nine nonamer pairs identified according to the selection criteria (Table S6, figure 4A). The VHFFKNIVT nonamer binds to the HLAs DRB1*08:02 and DRB1*15:01, forming two pairs selected by minimum 1 criterion (P2-P3-P5-P8). It is remarkable for being present in 15-mer peptides immunogenic (Table 3). Another pair selected by criterion 1 (P2-P3-P5-P8) was FKNIVTPRT^MBP/YKNRVASRK^EBV, both HLA-DRB5*01:01 ligands. Other nonamers also classified as possibly immunogenic are FKNIVTPRT and IRLFSRDAP.

#### Periaxin (PRX)

PRX presented thirty-five pairs of nonamers (Table S10, figure 4I). HLA-DQA1*05:01/DQB1*03:01 has the highest number of nonamer pairs (22). The VGVDLALPG nonamer forms three pairs; one of them, with the FGSDLALPS nonamer of the BOLF1 protein, presented a similarity score of 29 and greater identity (6/9). Another pair with a similarity score of 29 was (SETGAPGPA^Periaxin/ENTGAPAPP^EBV) and identity 5/9. The nonamer PRPAAPEVV is present in the highest number of pairs, 6. A pair formed by the nonamer TEAAELVPG was selected by criterion 1 (P2-P3-P5-P8). The YEDALRLQ nonamer stands out for participating in 15-mer peptides predicted to be immunogenic (Table 3). Four pairs bind to HLA-DQA1*01:02-DQB1*06:02.

#### Proteins predicted as minor cross-reactivity chance

CNP, MAG, MOBP, P0, P2, and PLP proteins are less likely to cross-react due to the low number of predicted pairs, absence of immunogenic peptides, or the lack of peptides binding to multiple sclerosis-risk HLA. In CNP, eight pairs of nonamers were selected, all by criterion 2 (P5 - 2 x P2/P3/P8 - 1 x P4/P6/P7) (Table S3, figure 4C). The nonamer TTGARVELS and VEAVQTGLD are part of the two pairs with the highest identities (5/9). Furthermore, two pairs (YKITPGARG^CNP/LKIHPGVAM^EBV, TTGARVELS^CNP/AEGLRVLLA^EBV) bind to HLAs of the DR15 haplotype. Eight pairs of MAG protein nonamers with relevant identity to EBV nonamers were found, all selected by criterion 2 (P5 - 2 x P2/P3/P8 - 1 x P4/P6/P7) (Table S5, figure 4E). The nonamer AESLLLELE is notable by forming three pairs, one of which has the highest similarity value (22). The MOBP protein has only one pair of nonamers with four identical residues and binds to HLA-DRB5*01:01 (Table S7, figure 4F). Five pairs of nonamers were selected from the P0 protein, all classified by criterion 2 (P5 - 2 x P2/P3/P8 - 1 x P4/P6/P7), and have four identical residues (Table S8, figure 4G). The most relevant nonamers were LVLSPAQAI, which forms the pair with the most remarkable similarity (21), and VLGAVIGGV, for being in two pairs. Four pairs of nonamers for the P2 protein were found, all selected by criterion 2 (P5 - 2 x P2/P3/P8 - 1 x P4/P6/P7), and had four identical residues (Table S9, figure 4H). The nonamer VGLATRKLG is prominent because it occurs in two pairs. We found eight pairs of nonamers for the PLP protein (Table S11, figure 4J). Nonamers containing the segment (F)VGAAATLV(S) are present in five pairs, all with the participation of HLA-DQA1*05:01/DQB1*03:01, and four of them with five identical residues. The FYTTGAVRQ nonamer binds to three different HLAs, forming three pairs. One nonamer (FVGAAATLV) is present in 15-mer peptides predicted to be immunogenic (Table 3).

## Discussion

In this study, a broad cross-identity prediction of essential residues was performed aiming to identify the interaction with the TCR between pairs of EBV and nervous system protein nonamers. All ten nervous system proteins presented nonamers pairs and are, thus, possible targets for self-reactive CD4+ T-cells. Some nervous system proteins also exhibited immunogenic peptides, suggesting more significant potential to stimulate CD4+ T cells. Prediction of peptides by the two methods produced 1695 results for nervous system proteins and 24912 for EBV proteins. This strategy has a limitation, as potential HLA-binding peptides predicted by only one server are not selected and could also be relevant. However, we chose to work with two methods based on the logical principle that a peptide is more likely to be a ligand if both approaches predict it.

Our result for the search for central nervous system proteins is consistent with proteome analyses that reported CNP, MAG, MBP, MOBP, and PLP among the highest relative abundances in the CNS (38). These proteins are prominent candidates in MS pathology because they are expressed in greater abundance and, to a greater extent, in CNS myelin than in other tissues. Proteome analyses found P0 and Periaxin to be the most abundant proteins in rat PNS myelin (35,36).

We investigated nervous system proteins regardless of their cellular location. Peptides present by MHC II molecules originate from the plasmatic membrane and the nuclear, intracellular, and extracellular compartments (39,40). The capture of antigens by antigen-presenting cells (APCs) can occur through a series of mechanisms, such as macropinocytosis, endocytosis, and phagocytosis (41). These three antigen capture mechanisms pick up extracellular material and transport it into the cell. This premise allows us to infer that MHC Class II molecules can present proteins from any cell compartment if they leak into the extracellular environment at some point. In healthy samples, HLA ligands are primarily membrane and extracellular proteins (42). However, presenting MHC II antigens must be considered in the context of inflammatory processes, such as what happens in autoimmune diseases. Through the mechanism known as epitope spreading, after initial tissue damage, cellular contents leak into the extracellular space, exposing autoantigens that can be captured by APCs and presented to self-reactive CD4 T-lymphocytes (43,44). It is likely that the epitope spreading mechanism initiates a cascade of self-reactivity years before the onset of clinical symptoms (45). A previous EBV infection likely triggers multiple sclerosis, but its role in disease progression is poorly understood (46,47).

The MBP protein has the highest absolute abundance in the CNS, but the relative abundance is the same as in the PNS (48). MAG and CNP proteins are also abundant in both myelin. Thus, one may question why self-reactive lymphocytes against these proteins, especially MBP, do not affect the PNS. Autoimmunity in MS may occur against antigens shared by peripheral and central myelin, such as MBP and MAG (49). Indeed, clinical symptoms and neurophysiological signs indicate that the PNS is affected in some patients with MS (49). However, multiple environmental and genetic factors trigger both MS and GBS. Thus, the mere sharing of antigenic structures, such as myelin proteins, or a recent infection by the same etiological agent, such as EBV, is not enough to explain the clinical condition developed by the patient.

Myelin protein P0, Myelin protein P2, and Periaxin are some of the most abundant proteins in the PNS (35,36). Studies examining P0 and P2 proteins in patients with GBS have been inconclusive regarding their role in pathogenesis. Thus, the low number of pairs identified and no peptide predicted to be immunogenic indicate that they have a low potential to exhibit molecular mimicry with EBV proteins and to be targets of autoimmunity. One of the first studies to analyze the CD4 T-lymphocyte response in patients with GBS showed that all peptides of P0 (56–71, 70–85, 180-199, and 194-208) and P2 (14-25 and 58-81) examined caused a proliferation of CD4 T-lymphocytes (50). However, in another work, only peptide P0 180-199 significantly stimulated CD4 T-lymphocyte proliferation compared with a healthy control group (51). Some studies showed that these peptides induced the secretion of cytokines, particularly IL-4 and TGF-β (52,53) – indicating the involvement of T-cells - but another study did not observe a similar result (51). The most comprehensive study evaluating the immune response in patients with GBS examined peptides corresponding to almost the entire amino acid sequence of the P0, P2, and PMP22 proteins (54). The authors found that IFN-γ production in response to the peptides was low or absent in most samples. Given these results, these proteins may not be a target of CD4+ T-cells in GBS. However, few papers have investigated peripheral myelin protein epitopes that might elicit a CD4 T-lymphocyte response. Therefore, additional epitopes not yet investigated from these or other proteins could be targets of T-cells. One example is Periaxin, which has never been studied as a source of autoantigens in GBS, although it is most abundant in peripheral myelin. However, our results showed several pairs that could cause molecular mimicry.

As observed in this work and demonstrated in the literature, Proteolipid protein (PLP) is the most abundant myelin protein in the CNS (38). Some studies have detected higher reactivity to PLP epitopes in MS patients than controls. However, others have not detected a difference in reactivity (55), similar to the case of MBP. MAG proteins, MOBP, and CNP, although less investigated than MBP and PLP, have also been reported to stimulate B and CD4 T-lymphocyte responses in MS (55). Recently, reports identified the GlialCAM protein as a potential target of molecular mimicry in MS (12). Although APLP1 is abundant in CNS myelin, we did not find any research that examined it as a possible source of autoantigens in MS. However, our results suggest that it may also be a target of autoimmunity.

The nine residues in the MHC binding groove constitute the TCR binding core, whereas the residues outside the groove are called flanking residues. Although the flanking residues may contribute to the TCR-pMHC complex stability and the TCR affinity for the epitope (56,57), they were not considered in this analysis because it is the anchoring residues that confer specificity to the TCR and interact with the complementarity determining regions (CDRs) (27,28,30,58). Cross-reactivity does not require that all residues are the same, but it is sufficient that some critical TCR contact residues are identical (25–30). Our strategy also considered the similarity between nonamers because similar peptides favor the occurrence of T-CD4 cross-reactivity, even if the identity is low (59).

The criteria we have defined consider the canonical diagonal orientation of the TCR-pMHC topology, in which the CDR3α and CDR3β loops are positioned over the central residue (P5) (60). However, unconventional binding topologies exist in which the CDR loops primarily contact other residues. One example is the TCR of clone Ob.1A12, which contacts residues only in the N-terminal half (P-4, P-2, P-1, P2, P3, and P5) of the MBP peptide. Also, in this clone, the sharing of the so-called "HF" motif (amino acids P2 His, P3 Phe, and P5 Lys/Arg) is necessary to stimulate T-cell proliferation (28), and positions P2 and P3 form the primary contacts with the TCR. Therefore, our proposed search strategy may not be efficient in detecting possible cross-reactions of TCR-pMHC complexes that do not obey conventional topology. Another limitation of our proposal is that it disregards the affinity of the T-cell clone for the nonamer. Populations of T-cells that have undergone an extensive clonal deletion process have low TCR affinity and are more sensitive to changes in MHC anchoring residues (27).

The P5 residue is probably the most important for TCR recognition because the CDR3 loops usually contact the amino acid in this position (60). Other evidence that supports the relevance of P5 is the observation that only the substitution of this amino acid inhibits the T-cell clone proliferative response and is the only conserved residue between two peptides recognized by the same TCR (25). Analysis of the structure of the TCR-pMHC complex also confirmed the importance of P5 (61,62). In an analysis of the contribution of each peptide residue to the reduction of T-cell reactivity, the substitution of P5 was more relevant, followed by substitutions in P2 or P8, in P4, P6, or P7 and P1, P9 or the flanking residues (27). The side chains of residues P2, P3, P5, and P8 generally point outside the binding groove and contact the TCR (27,63). P7 and possibly P3 bind the TCR and the MHC (27). P4 and P6 rarely interact with TCR, although with only a few atomic contacts, and can also affect its specificity indirectly by altering the conformation of other nonamer residues (27,58,62). Based on this information, we defined P5 as the main one, 2 and 8 as secondary and 3, 4, 6, and 7 as tertiary. Previous work has also explored molecular mimicry based on nonamer identity but has not considered the relevance of TCR contact residues (64,65). As far as we know, this is the first work to assess the potential for autoimmunity based on the identity of critical TCR contact residues.

Our investigation conclusively identified 133 pairs of nonamers that satisfy one or more of the selection criteria, indicating a high potential for these nonamers to be targeted by autoimmunity. 11 also contain immunogenic 15-mer peptides. Notably, various HLAs and haplotypes bind to nonamer pairs, indicating that different individuals - whose HLA haplotypes vary - may have autoimmunity through self-reactivity directed to different antigens, which bind to different HLA molecules. MS and GBS may have different autoantigens since their immunology was heterogeneous (45,66,67). In MS, the self-reactivity profile can vary even among patients (45,66), which suggests a diversity of immune targets relevant to its development without any particular antigen being responsible for the pathogenesis.

MBP is the most studied protein as an antigen in MS (68). There is long-standing evidence of recognition of MBP peptides containing the nonamer VHFFKNIVT by autoreactive CD4 T-lymphocytes in the context of HLA DR2 (DRB1*1501 and DRB5*0101) (69,70). The 18.5-kDa isoform of MBP is the most abundant in adult human myelin (68). The nonamers DSAATSESL and IRLFSRDAP, present in two selected pairs, are not contained in this isoform. Increased reactivity against the 82–102 region of MBP (numbered according to the 18.5 kDa isoform) (DENPVVHFFKNIVTPRTPPP) has been observed in patients with MS (71) and during the relapse phase of the disease (72). We find one pair with the nonamer VHFFKNIVT, which fits into this sequence. Other studies have found no difference between MS and control groups (73–75), consistent with the hypothesis that the immune response to MS is heterogeneous and varies from patient to patient. The MBP 85–105 peptide appears to be able to bind to the HLAs DRB1*1501 and DRB5*0101, which constitute the DR15 haplotype and are of particular interest in MS (76,77). One of the pairs found, consisting of the nonamers VHFFKNIVT (MBP) and YHFVKKHVH (BALF5), was described as being involved in molecular mimicry between MBP and EBV DNA polymerase (BALF5) (78). The Hy.2E11 T-cell clone isolated from MS patients recognizes the MBP 85–99 epitope restricted on HLA-DRB1*15:01 and the EBV627–641 presented by HLA-DRB5*01:01 (26). In this study, in addition to being predicted to bind to the same HLAs demonstrated by Lang et al., they also bind to HLA-DRB1*08:02. In a study investigating the immunoreactivity of antibodies and CD4 T-lymphocytes against MBP in patients with GBS, no difference was observed in the control group (51,79).

Previous studies have demonstrated that cross-reactivity does not necessarily require that the same HLA presents the peptides, but it is enough that the TCR contact residues are the same (25,26,30). Thus, we compare two nonamers predicted to bind to the same HLA or two different HLAs in the same haplotype. While the discovery of 1411 haplotypes may initially appear overwhelming, it is critical to note the significance of this number in the broader context of immunology. This extensive range of haplotypes not only confirms the diversity of immune responses but also suggests that the interactions between these HLAs and the identified proteins could be highly specific. Understanding this specificity could pave the way for targeted therapies.

Alleles comprising the DR15 haplotype (DRB1*15:01, DRB5*01:01, and DQA1*01:02/DQB1*06:02), known to be a risk factor for MS (37), were predicted to bind to pairs in this work. Genome-wide association studies have shown that HLA class II DRB1*15:01 has the strongest risk association with MS (80). This allele is also associated with the premature development of MS (81). Other MS-associated HLAs include DRB1*13:03, DRB1*03:01, DRB1*08:01, and DQA1*01:02/DQB1*03:02, which confers risk with different effect sizes (37,82). Studies of European-originated populations demonstrated an association of these alleles with risk. However, it is essential to study the genetics of MS in populations with diverse backgrounds to yield insights that may benefit individuals with MS from all ethnic groups (83). HLAs absent or uncommon in populations of European ancestry have been identified as risk factors in other ethnic groups, such as DRB1*04:05 and DRB1*15:03 (84). Predictions indicate that the DRB1*04:05 allele carries a GlialCam protein pair, which may be related to molecular mimicry. HLA-DRB1*15:01 has also been associated with risk in Jews, Sardinians, Africans, Hispanics, Japanese, and Indians (84). Furthermore, an allele’s protective or predisposing role depends on the ethnic group in which it is present (85), which emphasizes the importance of including a greater diversity of individuals in immunogenomics studies (83).

In a study carried out in the Tunisian population, the HLA-DRB1∗13:01 and DRB1∗14:01 alleles, as well as the DRB1*13/DQB1*03 and DRB1*14/DQB1*05 haplotypes, were found more frequently in patients with AIDP, while the prevalence of HLAs DRB1*03 and DRB1*07 was reduced (86). Chinese patients with AIDP have high frequencies of HLA DRB1∗13:01 (87). The DR16 and DQ5 alleles frequencies showed a significant increase in the AIDP group compared to the control (88). Another work reported elevations in the frequency of HLAs DRB1*03:01, DRB1*07:01, and DRB4*01:01 in Iraqi patients with the demyelinating form of GBS (89). A meta-analysis demonstrated no significant association between polymorphisms in HLA-DQB1 and risk for GBS in Asian and Caucasian populations (90). Although these works indicate a possible predisposing role of HLAs for AIDP, studies are still scarce and only performed in some specific populations. Therefore, it is still early to state that they are risk alleles for developing AIDP. In addition, studies do not always distinguish GBS subtypes, which makes it difficult to establish specific genetic risk factors for each form of the disease.

All data generated by this work resulted from *In silico* analyses. Although this approach effectively screens for potential autoantigens, mere identity at critical TCR contact residues does not imply molecular mimicry nor that such peptides are immunogenic. Furthermore, to search for potential etiological agents and autoantigens that trigger the autoimmune response, experimental research is essential to find HLAs associated with GBS and MS - mainly in non-European ethnicity populations. These data are scarce for Guillain-Barré syndrome, especially for the AIDP subtype. The pairs of identical nonamers found in this work provide evidence of cross-reactivity and can support the investigation of autoantigens in laboratory experiments.

## Conclusion

Our study is the first to extensively search for potential protein autoantigens in Guillain-Barré syndrome and multiple sclerosis, presented by common HLAs in the global population. We employed a novel approach based on sequence identity at critical TCR contact residues. The results revealed that several peptides derived from proteins in the nervous system and the Epstein-Barr virus share identity in these critical residues, suggesting the possibility of cross-reactivity between them. Notably, the Periaxin protein, not previously reported as a source of autoantigens, presented several nonamers with the potential to be related to molecular mimicry. Given its high expression in the peripheral nervous system (PNS), it could serve as an antigen in Guillain-Barré syndrome (GBS) or other peripheral neuropathies. Moreover, our study also examined CNS protein antigens (MBP, CNP, GlialCam, MAG, and PLP) that are already known targets of autoimmunity. We identified several possible nonamers within these CNS proteins that may contribute to molecular mimicry processes.

## List of abbreviations

GBS: Guillain-Barré Syndrome
MS: Multiple sclerosis
AIDP: Acute Inflammatory Demyelinating Polyradiculoneuropathy
PNS: Peripheral Nervous System
CNS: Central Nervous System
nTPM: normalized to transcripts per million
AFND: Allele Frequency Net Database
HLA: Human Leukocyte Antigen
TCR: T-cell receptor

## Acknowledgement

The authors would like to thank the Coordenação de Aperfeiçoamento de Pessoal de Nível Superior (CAPES) and Conselho Nacional de Desenvolvimento Científico e Tecnológico (CNPq) for their financial support. H.A. Patrocínio and T.S. Fiúza had a scholarship from CAPES (Finance code: 001). The authors are indebted to the High-Performance Computing Center (NPAD/UFRN) and the Multidisciplinary Bioinformatics Environment (BioME/UFRN) for providing computing resources.

## Data Availability

The code used for analyzing nonamer cross-identity, processing the data, and generating the figures is available at https://github.com/evoMOL-Lab/immuno-cross. Supplementary Material is available through the following [link](https://github.com/evoMOL-Lab/immuno-cross/tree/main/Supplementary_Material).

## Notes

https://github.com/evoMOL-Lab/immuno-cross

### Competing Interest Statement

The authors have declared no competing interest.

